# Environmental and genetic influence on rate and spectrum of spontaneous mutations in *Escherichia coli*

**DOI:** 10.1101/2023.04.06.535897

**Authors:** Danna R. Gifford, Anish Bhattacharyya, Alexandra Geim, Eleanor Marshall, Rok Krašovec, Christopher G. Knight

## Abstract

Spontaneous mutations are the ultimate source of novel genetic variation on which evolution operates. Although mutation rate is often discussed as a single parameter in evolution, it comprises multiple distinct types of changes at the level of DNA. Moreover, the rates of these distinct changes can be independently influenced by genomic background and environmental conditions. Using fluctuation tests, we characterised the spectrum of spontaneous mutations in *Escherichia coli* grown in low and high glucose environments. These conditions are known to affect the rate of spontaneous mutation in wild-type MG1655, but not in a Δ*luxS* deletant strain —a gene with roles in both quorum sensing and the recycling of methylation products used in *Escherichia coli*’s DNA repair process. We find an increase in AT>GC transitions in the low glucose environment, suggesting that processes relating to the production or repair of this mutation could drive the response of overall mutation rate to glucose concentration. Interestingly, this increase in AT>GC transitions is maintained by the glucose non-responsive Δ*luxS* deletant. Instead, an elevated rate of GC>TA transversions, more common in a high glucose environment, leads to a net non-responsiveness of overall mutation rate for this strain. Our results show how relatively subtle changes, such as the concentration of a carbon substrate or loss of a regulatory gene, can substantially influence the amount and nature of genetic variation available to selection.

## Introduction

Understanding what governs the occurrence of spontaneous mutations is crucial for predicting evolution[1–3]. Mutation rate is known to be influenced by genetic variation [4, 5] and the environment [6–8] (i.e. mutation rate plasticity). Although often described as a single metric, mutation rate encompasses the cumulative effects of numerous biochemical processes that cause errors in DNA replication to occur and remain uncorrected. Progress in understanding the molecular basis for mutation has been made by characterising the ‘mutational spectrum’, i.e. the frequency of occurrence of different types of single nucleotide variants (SNVs) and insertion and deletion mutations (indels). For SNVs, this includes a set of six possible mutations, the transitions AT>GC and GC>AT, and the transversions AT>CG, AT>TA, GC>CG, and GC>TA. For indels, the set of possible mutations is much greater [9]. Each type of SNV and indel occurs at different rates, which can each respond differently to genetic and environmental factors, [7, 8, 10–15].

In microbes, both the rate and spectrum of spontaneous mutations change plastically with aspects such as nutrient availability [12, 16, 17], growth rate variation[7], and temperature[8, 16, 18], providing insight into how mutation generating processes operate at a molecular level.

Observing changes in the mutational spectrum in deletion strains has also been used to characterise the roles of genes involved in mutation avoidance and repair systems. For instance, disruption of MMR increases the frequency of the transitions, AT>GC and GC>AT[19]. Disruption of guanine oxidation (GO) system genes *mutM* or *mutY*, which typically correct errors induced by oxidative damage, increases the frequency of GC>TA transversions [11], and disruption of *mutT* increases AT>CG transversions [10, 20]. Differential expression of different DNA polymerase genes can also produce striking effects on the mutational spectrum [21]. Collectively, these studies provide insight into how mutation-generating processes operate at a molecular level.

We have previously described an inverse relationship between the rate of spontaneous mutation and glucose concentration [22], or, more generally, nutrient availability [23], in *Escherichia coli*. This association is detectable across the diversity of *E. coli* [24], and is also present in other bacteria, in archaea, and in yeast [24, 25]. The strength of the association between glucose concentration and overall mutation rate varies within organisms from the same species [24, 25] and can be either enhanced or abolished by certain genetic knockouts [22, 23, 25]. Despite the pervasiveness of this phenomenon, we do not currently know whether it arises due to an overall increase with in all possible types of mutation, or whether particular classes of SNVs or indels are increased. Previous results indicate that mutation avoidance and repair systems and metabolic effects seem to play a part in mutation rate plasticity. For instance, the association between glucose and overall mutation rate is diminished in strains with impaired mutation avoidance and repair (through deletion of *mutS, mutH, mutL* or *mutT*[25]). Yet, not all genes affecting mutagenesis and repair affect mutation rate plasticity; deletion of polymerase IV or polymerase V, for example, does not have this effect [23, 25]. This shows that the molecular basis for the mutation rate response to glucose may be connected to interactions between metabolism and mutation avoidance and repair.

Intriguingly, the association between glucose concentration and overall mutation rate was also greatly diminished in a strain with a deletion of the *luxS* gene [22], which is involved in both AI-2 mediated quorum sensing and the activated methyl cycle (AMC) [26]. This raises the question of whether the response of the overall mutation rate to glucose concentration in the wild-type MG1655 strain is due to population-density effects mediated by quorum sensing, or via an interaction between the AMC and DNA replication fidelity. The latter possibility arises due to the role of the AMC in generating the methyl-group donor *S*-adenosyl-*L*-methionine (SAM), which is required for DNA methylation and *E. coli*-specific methyl-directed mismatch repair (MMR) [27].

Thus, *luxS* could potentially be deficient in methyl-directed MMR. Alternatively, the effect of *luxS* on overall mutation rate may emerge due to altered gene regulation more generally. Deletion of *luxS* influences gene expression in an environment-dependent manner [28]. In particular, *luxS* affects regulation of metabolism, nutrient acquisition and virulence traits [29, 30], which could influence how this strain responds to and metabolises glucose.

Observing how the spectrum of spontaneous mutations shifts in response to glucose concentration and deletion of *luxS* can provide insight into some of the molecular processes that underpin the response of mutation rate to glucose in *E. coli*. To accomplish this, we have simultaneously measured the response of mutation rate to glucose via fluctuation tests, and characterised how the spectrum of spontaneous mutations differs in high and low glucose environments. Our results are consistent with a previous finding that lower overall mutation rates were observed under high glucose environments in wild-type *E. coli*, but not a Δ*luxS* deletion strain [22]. Curiously, however, both strains exhibited a decrease in AT>GC transitions under high glucose conditions (a finding consistent with ref. [16]). This indicates that the deletion of the *luxS* gene does not appear to directly diminish the connection between glucose and mutation rates, as seen in the wild-type strain. Instead, the Δ*luxS* deletant demonstrated a higher occurrence of a different single nucleotide variant (SNV), specifically GC>TA transversions. For Δ*luxS*, these GC>TA transversions were observed under both glucose concentrations, but were more common under high glucose conditions. This makes the Δ*luxS* strain appear to be non-responsive to glucose when, in reality, separate mechanisms increase and decrease particular elements of the mutational spectrum as glucose concentration changes, in a way that cancels out when these separate elements are combined into a single overall mutation rate. Together this supports the idea that seemingly small-scale genomic and environmental changes can have a significant effect on the quantity and nature of genetic variation available to selection.

## Materials and methods Strains and media

Experiments used *E. coli* str. K-12 substr. MG1655 as the wild-type and a Δ*luxS* deletant (KX1228), which was constructed in the same genetic background [31]. We used lysogeny broth for routine culturing of strains [LB, 10 g/l tryptone (ThermoFisher Scientific, UK), 5 g/l Bacto yeast extract (BD Biosciences, UK), 10 g/l NaCl (ThermoFisher Scientific, UK)]. Where indicated, LB agar was prepared by adding 12 g/l agar (BD Biosciences, UK).

For fluctuation tests, we used Davis minimal medium [DM, 7 g/l potassium phosphate dibasic trihydrate, 2 g/l potassium phosphate monobasic anhydrous, 1 g/l ammonium sulfate, 0.5 g/l sodium citrate (all obtained from Sigma-Aldrich, UK)]. After autoclaving, we added filter sterilised magnesium sulfate (in each litre of medium, 500 of 10% w/v, Fisher Scientific) and thiamine hydrochloride (500 of 0.2% w/v, Sigma Aldrich, UK). We added D-glucose (ThermoFisher Scientific, UK) to DM as a carbon source, at concentrations of 80 mg/l (referred to as the ‘low glucose’ treatment) or 250 mg/l (referred to as the ‘high glucose’ treatment).

These specific concentrations were chosen based on their previously observed capacity to induce a discernible difference in mutation rates [22].

For plating of the fluctuation tests, we used tetrazolium agar (TA, 10 g/l tryptone, 1 g/l Bacto yeast extract, 3.75 g/l NaCl, 12 g/l agar) with L(+)-arabinose (3 g/l, Sigma-Aldrich, UK) and 2,3,5-triphenyltetrazolium chloride (50 mg/l, Sigma-Aldrich, UK) added post-autoclaving. Selective media was TA supplemented with 50 mg/l rifampicin (ThermoFisher Scientific, UK), which was dissolved in 99% methanol (ThermoFisher Scientific, UK) before adding to warm (∼55 °C) medium.

### Estimating mutation rates via fluctuation tests

We estimated overall mutation rates using a high-throughput variation [32] of the classic fluctuation test [33, 34]. Briefly, overnight cultures of MG1655 and Δ*luxS* were initiated by inoculating single colonies (streaked on LB agar) into 5 ml LB in 50 ml Falcon conical centrifuge tubes (Corning, USA). Overnight cultures were diluted in DM to an optical density of 0.3 (OD, 600 nm), then subsequently diluted by a factor of 5×10^−4^ to achieve a density of 1 × 10^3^ to 5 × 10^3^bacterial cells per ml. From these dilutions, 0.5 ml was used to establish 182 independent cultures in 96-well deep well plates (Greiner Bio-One, Austria). Cultures grew at 37 °C with 200 RPM shaking for 24 h. Each independent culture was subsequently plated on 5 ml selective media, using 6-well plates (Greiner Bio-One, Austria) as the culture vessel. After 48 h growth, we counted the number of rifampicin resistant mutants that arose in each parallel culture. In concert, we estimated the population density of a subset of three parallel cultures, by plating serial dilutions on TA agar in 90 mm Petri dishes. We isolated resistant mutants from the selective plates and inoculated these into 1 ml cultures of LB for overnight growth, prior to storage in 75% LB and 25% glycerol at −80 °C in 96-well polypropylene plates (ThermoFisher Scientific, UK). To ensure mutations were independent, only one colony was isolated from independent cultures where mutants appeared (and nothing was isolated from cultures where no resistant mutants appeared).

### Determining the spectrum of spontaneous resistance mutations

We used resistant mutants arising from the fluctuation tests to analyse the spectrum of spontaneous mutations. As our hypothesis about mutation rate plasticity involved changes in SNVs, our study required a selective marker that would reveal a diverse spectrum of SNVs. Rifampicin resistance, extensively used for characterizing mutational spectra, is well-suited to this purpose [35–38]. More than 80 distinct SNVs associated with rifampicin resistance have been identified within the *rpoB* gene. These mutations predominantly occur within the rifampicin resistance determining region (RRDR) of *rpoB*, consisting of three clusters: cluster I (amino acid positions 507-533/nucleotide positions 1520–1598), cluster II (amino acid positions 563–572/nucleotide positions 1687–1715), and cluster III (amino acid position 687/nucleotide positions 2060-2062). As *rpoB* is an essential gene, loss of function mutations cannot confer resistance, making it an effective marker for observing SNV changes over indels. Although in-frame indels conferring resistance do occur occasionally, they are proportionally fewer compared to SNVs. In contrast, antibiotics where resistance arises due to loss of function mutations typically exhibit higher proportions of indels, along with insertion sequence disruption, as seen in D-cycloserine resistance via the *cycA* gene [39].

For each isolated mutant, we amplified a region of the *rpoB* gene containing clusters I and II of the RRDR, which could be captured via a single PCR using forward primer 5’-ATGATATCGACCACCTCGG-3’ and reverse primer 3’-TTCACCCGGATACATCTCG-5 (Integrated DNA Technologies, Belgium). Primers were resuspended and diluted to a stock concentration of 100 M in nuclease-free water prior to use. To obtain DNA for Sanger sequencing, resistant mutants were revived from the freezer by inoculating 1 of culture on to LB agar using a 96-well pin replicator (Boekel Scientific, USA). Following overnight growth, a pipette tip was used to scrape a small amount of colony material into 10 nuclease-free water.

Each of 1 forward primer stock, 1 reverse primer stock, and 12.5 master mix (Platinum Green Hot Start PCR Master Mix, ThermoFisher Scientific, UK) was added to each reaction tube, and was mixed gently by pipetting. PCRs were performed as follows: i) initial denaturation (94 °C for 5 min), ii) equilibration (30 °C for 5 min), iii) denaturation (94 °C for 0.5 min), iv) annealing (62 °C for 0.5 min), v) extension (72 °C for 0.5 min), vi) repeat steps iii–v for 35 cycles, vii) final extension (72 °C for 10 min), viii) hold at 4 °C. PCR products were purified using the QIAquick PCR Purification Kit (Qiagen, UK). Purified DNA was diluted to approximately 5 ng/. Sanger sequencing was performed by Eurofins LightRun service (Eurofins Genomics, Germany) using the reverse primer indicated above as the sequencing primer.

Mutations were identified by aligning Sanger sequence data to the *rpoB* of *E. coli* str. K-12 [NCBI accession NC_000913.3 (4181245..4185273)] using Unipro UGENE version 33 obtained from http://ugene.net [40] (data provided in File S1). As we acquired different numbers of mutants from the fluctuation tests, we present SNVs in as frequencies rather than absolute counts.

## Statistical analysis

### Data processing and mutation rate estimation

Data processing and statistical analyses were performed in R (version 4.0.3) [41] using functions from the tidyverse [42] (analysis code provided in File S2). Figures were produced using ggplot2 [43]. We used `mutestim()` (with default settings) from the flan package (version 0.9) [44] to estimate mutation rates from the mutant counts and population sizes obtained via fluctuation tests (data provided in Files S3 and S4).

### Mutation rate comparison

We compared ratios of overall mutation rates measured at different glucose concentrations that previously elicited a difference in mutation rates in wild-type MG1655, but not the Δ*luxS* deletant (while not affecting growth rate) [22]. We calculated a low-to-high ratio for mutation rates, i.e. a ratio close to 1 indicates a limited response to glucose, a ratio > 1 indicates a higher mutation rate at low glucose, and a ratio < 1 indicates a lower mutation rate at low glucose. We first calculated point estimates of the mean and variance of these ratios (the latter using the ‘sigma method’ [45]), and then performed a *z*-test on the ratio for each strain (for full details, see supplemental material).

### Mutational spectrum comparison

We assessed the effect of glucose concentration and strain on the overall mutational spectrum by performing multinomial logistic regression on counts of each SNV observed, using `multinom()` from the nnet package [46]. As their numbers were small, all insertions were grouped into one category, and all deletions into another category. We subsequently looked at individual SNV categories by fitting logistic regression models on each SNV separately using `glm()` with a binomial family. We used Type II ANOVA (with a likelihood-ratio *χ*^2^ as the test statistic) to determine the significance of removing each predictor variable in the presence of other predictor variables using the `Anova()` function from the car package [47]. We tested only a single planned comparison within a model, therefore *p*-values are given directly from this ANOVA without further correction [48].

## Results and discussion

A shows that wild-type MG1655 and Δ*luxS* deletant have very different mutation rate responses to glucose concentrations in minimal media. To assess the change in mutation rate with environment, we calculated the ratio of mutation rates estimated in the low and high glucose concentration treatments, which we refer to as the ‘low-to-high ratio’. We observed a significantly higher mutation rate at low glucose in the wild-type MG1655 [low-to-high ratio: 1.60, 95% CI = 1.58,1.62], but only a marginal change in Δ*luxS* [low-to-high ratio: 1.06, 95% CI = 1.00, 1.12], which is consistent with our previous research [22–25]. The wild-type MG1655 and Δ*luxS* low-to-high ratios were significantly different (*z*-test: *Z* = 2.69, *p* = 0.004).

To characterise the spectrum of mutations underlying these differences in mutation rate, we sequenced the *rpoB* gene of resistant isolates that arose under low and high glucose conditions (B). Mutations were found in the region of *rpoB* sequenced (RRDR clusters I and II) for 214 out of 274 sequenced isolates (Table S1 and File S1); isolates where no mutation was detected may have had mutations in another portion of *rpoB* [49], or a resistance mechanism not involving *rpoB*. The majority of mutations observed were SNVs at loci previously described to give rise to rifampicin resistance [21, 35–37]. We also observed 10 isolates with in-frame insertions or deletions, which were significantly more frequent in the wild-type MG1655 genetic background, but not significantly affected by glucose concentration (logistic model, glucose concentration: *χ*_1_^2^ = 0.01, *p* = 0.89; strain: *χ*_1_^2^ = 5.47, *p* = 0.019; interaction: *χ*_1_^2^ = 0.81, *p* = 0.36). We observed no indels that would result in frame shifts, which is expected given that *rpoB* is an essential gene [50].

Across all types of mutations, we found a significant effect of strain, but no significant effect of glucose concentration, or their interaction on the mutational spectrum as a whole (multinomial regression model, glucose concentration: *χ*_7_^2^ = 11.9, *p* = 0.10; strain: *χ*_7_^2^ = 15.2, *p* = 0.034; interaction: *χ*_7_^2^ = 5.92, *p* = 0.55). Subsequently, we examined the impact of genotype and environment on the frequency of individual mutational categories. For SNVs, we observed distinct shifts in two classes, AT>GC transitions and GC>TA transversions. AT>GC transitions were significantly higher in the low glucose environment, but were unaffected by strain (logistic model, glucose concentration: *χ*_1_^2^ = 5.44, *p* = 0.020; strain: *χ*_1_ ^2^= 0.17, *p* = 0.68; interaction *χ*_1_^2^ = 0.67, *p* = 0.41). A shift toward AT>GC transitions under glucose-limited conditions was previously observed for *E. coli* str. MC4100 by Maharjan and Ferenci[16] using a chemostat system and a different selective marker (D-cycloserine resistance arising from loss of function mutations in *cycA*), suggesting the observation of elevated AT>GC transitions is not a result of our experimental conditions.

In contrast, GC>TA transversions were significantly elevated in the Δ*luxS* deletant, with a slight but non-significant increase in both strains in the high glucose treatment and no significant interaction (logistic model, glucose concentration: *χ*_1_^2^ = 3.45, *p* = 0.063; strain: *χ*_1_^2^= 5.21, *p* = 0.022; interaction: *χ*_1_^2^ = 0.50, *p* = 0.48). For all other SNVs, we did not observe a significant effect of either glucose concentration or strain (logistic models, all p>0.1). Some SNVs were infrequently observed, which may limit our ability to detect differences in frequency between treatments. However, note that we did observe a significant difference for indels (*n* = 10), which were less frequent than all classes of SNVs (*n* ≥ 24) excepting GC>CG transversions (*n* = 8).

The increased mutation rate associated with growth in a low glucose environment, observed here (A) and previously[22–25], seems to arise due to an increase in AT>GC transitions B. This points to the potential involvement of two molecular processes that could be linked to the mutation rate response to glucose. Elevated rates of AT>GC transitions are associated with deficient MMR, which typically repairs both types of transition mutations, AT>GC and GC>AT. Consistent with this finding, we previously observed that deletion of key MMR genes diminishes the response of mutation rate to glucose [25]. Although we did not detect a significant increase in GC>AT transitions in low glucose (in fact, GC>AT transitions were overall less frequent in low glucose), the relative proportion of this class of SNVs is much smaller overall, making changes harder to detect. However, an elevated rate of AT>GC transitions, without a similar elevation of GC>AT transitions, has also been observed under conditions inducing thymine starvation in *E. coli* and other organisms. This could point toward a common mechanism involving cellular pyrimidine availability [51].

The apparent loss of response to glucose in a Δ*luxS* deletant (A) was not due to the abolition of the effect on AT>GC transitions, as this SNV behaved similarly between environments for both strains (B). This is contrary to our proposed hypothesis that the disruption of SAM production in Δ*luxS* deletants [52] causes a decrease in the efficiency of methyl-directed MMR (usually associated with an increase in AT>GC and GC>AT transitions). Instead, we observed increased frequencies of GC>TA transversions in the Δ*luxS* strain, making it seem that the responsiveness of mutations to glucose is abolished, although the increase in AT>GC transitions under low glucose conditions remains. Under high glucose, this increase in GC>TA transversions coincidentally roughly offsets the decrease in AT>GC transitions—this does not imply direct antagonism between these SNV categories, which have distinct mechanistic bases. Elevated GC>TA transversions are typically associated with oxidative damage, and are usually corrected by the GO system involving MutM and MutY. That similar effects are observed for Δ*luxS* raises the possibility that *luxS* has a yet-uncharacterised role in regulating either the GO system or the oxidative stress response in *E. coli*. Several other lines of evidence also suggest this possibility. In *Streptococcus mutans, luxS* is involved in regulating genes involved in recombination and base excision repair, namely *recA*, and the apurinic/apyrimidinic AP endonucleases *smnA* and *nth*, as well as oxidative stress response genes [53, 54]. In *E. coli*, similar AP endonucleases are required for filling in nucleotide gaps caused by the GO system [55]. Deletion of *E. coli luxS* also affects expression of multiple genes [56], including several involved in oxidative damage response [28], carbon utilisation, and biofilm formation [57].

We note potential limitations in our approach to characterising the mutational spectrum. We used rifampicin as a selective marker as an efficient means of detecting SNVs using Sanger sequencing. Using alternative selective markers could unveil different patterns in how the mutational spectrum responds to environment and genomic changes. In particular, our ability to detect insertions and deletions (indels), insertion sequence disruption, and other disruptive mutations is constrained because rifampicin targets *rpoB*, an essential gene. Using a selective marker involving loss-of-function resistance mutations would enable characterisation of these other types of mutational events [7, 16–18, 58]. Alternative methods utilising high-throughput technologies could also enhance throughput to uncover a wider range of mutations compared to Sanger sequencing approaches. (e.g. maximum-depth sequencing of bulk cultures [59] or pooled sequencing [58]). However, irrespective of selective marker or sequencing approach used, fluctuation tests can also only characterise mutations within defined genes. Employing a mutation accumulation and whole-genome sequencing strategy could provide a broader perspective on mutations across the entire genome [10, 11, 60], but such approaches are labour intensive and therefore less practical for studying multiple genotypes and environments.

Our results indicate that mutation rate plasticity associated with growth in different glucose concentrations in wild-type *Escherichia coli* results from changes in the rate of a specific mutational class, AT>GC transitions, and not a universal change in mutation rate for all classes of mutation. The changes in AT>GC transitions persist in a Δ*luxS* strain, which suggests that the increase in AT>GC is not associated with *luxS*-mediated quorum sensing. Moreover, the apparent reduction in the plasticity of overall mutation rate in Δ*luxS* is due to an increase in a separate mutational class, GC>TA transversions, which are more common under high glucose conditions. The net effect means that we observe a decrease in overall mutation rate for wild-type MG1655 under high glucose, but no such decrease for Δ*luxS*. Consequently, the association between nutrient availability, population density and overall mutation rate previously observed for many microbes [22–25] is not likely driven by quorum sensing [61], but instead may be shaped by mutagenesis or deficiencies in mutation repair, potentially linked with low resource availability. As the mutational spectrum can have downstream consequences for adaptive evolution [1–3, 62], understanding how population growth and density affect mutation generation, avoidance, and repair processes will be essential for understanding adaptive potential.

## Data availability

Strains are available upon request. File S1 contains the mutations found in sequenced rifampicin-resistant strains originating from fluctuation test and used to assess changes in mutational spectrum. File S2 contains the R analysis code used to perform all statistical analyses and generate figures. File S3 contains mutant counts used to estimate mutation rates to rifampicin resistance for MG1655 and Δ*luxS* strains grown at low and high glucose in the fluctuation test. File S4 contains population density data (*N*_*t*_) for MG1655 and Δ*luxS* strains grown at low and high glucose in the fluctuation test.

## Acknowledgments

The authors thank Karina Xavier (Instituto Gulbenkian de Ciência) for the gift of the Δ*luxS* deletant strain KX1228 and its MG1655 parent strain.

## Figure

**Figure 1:**
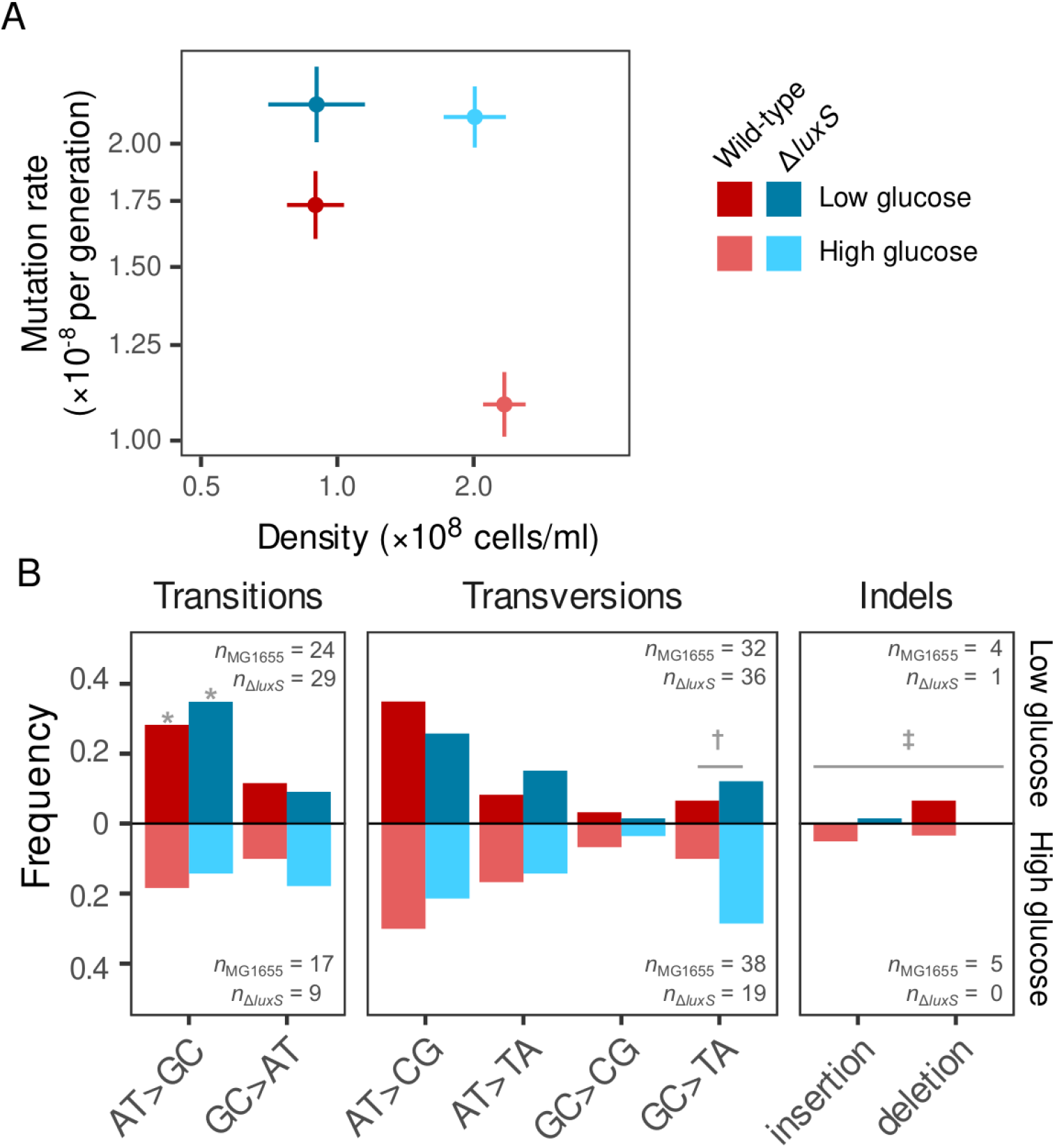
Effect of glucose concentration on rate and spectrum of mutations in *E. coli* str. K-12 MG1655 wild-type and Δ*luxS* deletant. A) Spontaneous mutation rate measured by fluctuation test (mean ± SE from *n* = 182 independent populations for each estimate). B) Spectrum of spontaneous mutations in wild-type MG1655 and Δ*luxS* deletant under high and low glucose treatments, showing increased AT>GC transitions in low glucose for both strains 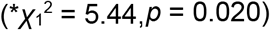, elevated GC>TA transversions for the Δ*luxS* deletant 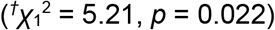, and a greater frequency of indels in the wild-type MG1655 background 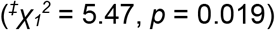. Note that relative frequencies are shown due to uneven sample sizes among treatments.

