## Supplementary Text for "Environmental and genetic influence on rate and spectrum of spontaneous mutations in *Escherichia coli*"

### Statistical analysis

#### Mutation rate comparison

The `mutestim()` function from the flan package[2,3] calculates point estimates for the mean ($\hat{u}$) and variance ($\widehat{\sigma^2}$) of mutation rates. To estimate for the response of mutation rate to glucose concentration, we calculated a point estimate for the ratio of mutation rates between low and high glucose conditions, which we define as the low-to-high ratio,

$$\hat{u}_{i\cdot\text{ratio}} = \frac{\hat{u}_{iLG}}{\hat{u}_{iHG}}, \tag{1}$$

where $LG$ and $HG$ represent low and high glucose conditions, respectively, and $i$ refers to strain, i.e. either wild-type or Δ*luxS*. This ratio indicates whether or not the mutation rate changes with glucose concentration. $\hat{u}_{i\cdot\text{ratio}} \approx 1$ indicates a limited response to glucose, $\hat{u}_{i\cdot\text{ratio}} > 1$ indicates a higher mutation rate at low density, and $\hat{u}_{i\cdot\text{ratio}} < 1$ indicates a lower mutation rate at low density.

We calculated a point estimate for the variance of this ratio using the sigma method[1],

$$\widehat{\sigma^2}[u_{i\cdot\text{ratio}}] = \frac{\text{Var}\,[u_{iLG}]}{\mathbb{E}[u_{iHG}]^2} - \frac{2\,\mathbb{E}[u_{iLG}]}{\mathbb{E}[u_{iHG}]^3}\text{cov}[u_{iLG}, u_{iHG}] + \frac{\mathbb{E}[u_{iLG}]^2}{\mathbb{E}[u_{iHG}]^4}\,\text{Var}[u_{iHG}]\,. \tag{2}$$

As $u_{iLG}$ and $u_{iHG}$ are independent estimates, $\text{cov}[u_{iLG}, u_{iHG}] = 0$.

We used the point estimates calculated by flan as estimators for the expectations and variances, which substituting into Equation 2 gives

$$\widehat{\sigma^2}[u_{i\cdot\text{ratio}}] = \frac{1}{\hat{u}^2_{iHG}}\left[\widehat{\sigma^2}\,[u_{iLG}] + \frac{\hat{u}^2_{iLG}}{\hat{u}^2_{iHG}}\,\widehat{\sigma^2}[u_{iHG}]\right]\,. \tag{3}$$

We then compared the ratios using a $z$-test on the difference in ratios between wild-type and Δ*luxS*, using the point estimates calculated in Equations 1 and 3 as estimators for the population statistics,

$$Z = \frac{\hat{u}_{\text{wild-type}\cdot\text{ratio}} - \hat{u}_{\Delta luxS\cdot\text{ratio}}}{\sqrt{\widehat{\sigma^2}_{u_{\text{wild-type}\cdot\text{ratio}}} + \widehat{\sigma^2}_{u_{\Delta luxS\cdot\text{ratio}}}}}\,. \tag{4}$$

A $p$-value was then calculated in R 4.0.3[4] as $1-$ `pnorm`$(Z)$.

**Table S1: Mutations in rifampicin resistant isolates found by Sanger sequencing of the *rpoB* rifampicin resistance determining region.** Mutations and their respective positions in coding sequence and protein sequence are given relative to the reference sequence for *E. coli* str. K-12 [NCBI Genome accession NC_000913.3 (4181245..4185273), NCBI Protein accession NP_418414.1]. Amino acid insertions indicate that the new amino acid has been inserted before the reference amino acid in the indicated position. A total of 214 mutations were found in 274 sequenced strains. Data also provided in File S1.

| Nucleotide position | Mutation | Coding change | MG1655 Low glucose | MG1655 High glucose | Δ*luxS* Low glucose | Δ*luxS* High glucose |
|---|---|---|---|---|---|---|
| 1525 | AT>CG | S509R | 0 | 1 | 0 | 0 |
| 1527 | GC>CG | S509R | 1 | 0 | 0 | 0 |
| 1527 | GC>TA | S509R | 0 | 1 | 1 | 1 |
| 1529-1531 | +GCT | ins511L | 0 | 1 | 0 | 0 |
| 1532 | AT>GC | L511P | 1 | 1 | 3 | 0 |
| 1532 | AT>CG | L511R | 3 | 1 | 0 | 1 |
| 1532 | AT>TA | L511Q | 1 | 2 | 1 | 1 |
| 1534 | AT>GC | S512P | 4 | 6 | 5 | 0 |
| 1535 | GC>AT | S512F | 1 | 0 | 1 | 0 |
| 1535 | GC>TA | S512Y | 0 | 0 | 1 | 1 |
| 1538 | AT>GC | Q513R | 1 | 0 | 1 | 0 |
| 1538 | AT>TA | Q513L | 0 | 1 | 1 | 0 |
| 1541-1546 | +TATGGC | ins514AM | 0 | 1 | 0 | 0 |
| 1542-1550 | -TATGGACCA | F514LdelMDQ | 1 | 0 | 0 | 0 |
| 1546 | GC>AT | D516N | 1 | 1 | 1 | 0 |
| 1547 | AT>GC | D516G | 8 | 4 | 8 | 3 |
| 1565 | GC>AT | S522F | 0 | 1 | 0 | 0 |
| 1574 | GC>CG | T525R | 1 | 0 | 0 | 0 |
| 1576 | GC>TA | H526N | 0 | 0 | 0 | 1 |
| 1577 | AT>TA | H526L | 0 | 1 | 0 | 0 |
| 1577 | AT>CG | H526P | 0 | 0 | 0 | 1 |
| 1578 | GC>CG | H526Q | 0 | 1 | 0 | 0 |
| 1578 | GC>TA | H526Q | 0 | 0 | 1 | 1 |
| 1585 | GC>AT | R529C | 1 | 0 | 1 | 0 |
| 1586 | GC>AT | R529H | 1 | 0 | 1 | 0 |
| 1589-1591 | +CTC | ins531S | 0 | 1 | 0 | 0 |
| 1590-1595 | -CTCCGC | del531SA | 1 | 0 | 0 | 0 |
| 1592 | GC>AT | S531F | 1 | 1 | 1 | 3 |
| 1592 | GC>TA | S531Y | 0 | 0 | 1 | 0 |
| 1593-1601 | -CGCACTCGG | de532ALG | 1 | 0 | 0 | 0 |
| 1595 | GC>TA | A532E | 0 | 0 | 1 | 0 |
| 1596-1598 | -ACT | del533L | 1 | 0 | 0 | 0 |
| 1598 | AT>GC | L533P | 1 | 0 | 6 | 1 |
| 1598 | AT>TA | L533H | 0 | 1 | 2 | 1 |
| 1600 | GC>AT | G534S | 1 | 1 | 1 | 0 |
| 1600-1608 | +CCGCACTCG | ins534AAL | 0 | 0 | 1 | 0 |
| 1601 | GC>TA | G534V | 2 | 1 | 0 | 1 |
| 1601 | GC>CG | G534A | 0 | 1 | 1 | 0 |
| 1603-1611 | -CCAGGCGGT | del535PGG | 0 | 2 | 0 | 0 |
| 1607 | GC>TA | G536V | 1 | 0 | 0 | 0 |
| 1610 | GC>AT | G537D | 0 | 0 | 0 | 1 |
| 1687 | AT>CG | T563P | 8 | 9 | 9 | 2 |
| 1691 | GC>AT | P564L | 0 | 1 | 0 | 0 |
| 1708 | GC>TA | G570C | 1 | 0 | 0 | 0 |
| 1709 | GC>CG | G570A | 0 | 2 | 0 | 1 |
| 1712 | AT>TA | L571Q | 1 | 0 | 0 | 0 |
| 1714 | AT>CG | I572L | 8 | 6 | 5 | 1 |
| 1714 | AT>TA | I572F | 1 | 2 | 2 | 0 |
| 1715 | AT>GC | I572T | 2 | 0 | 0 | 0 |
| 1715 | AT>CG | I572S | 2 | 1 | 3 | 1 |
| 1715 | AT>TA | I572N | 2 | 3 | 4 | 2 |
| 1721 | GC>AT | S574F | 1 | 1 | 0 | 1 |
| 1721 | GC>TA | S574Y | 0 | 4 | 3 | 3 |
| Mutations found/strains sequenced: | | | 60/82 | 60/77 | 66/79 | 28/36 |

[4] R Core Team, 2020 *R: A Language and Environment for Statistical Computing*. R Foundation for Statistical Computing, Vienna, Austria.
