## Supplementary material for "Environmental and genetic influence on rate and spectrum of spontaneous mutations in *Escherichia coli*": File S2 - R source.R

```
library(MASS)		# Load before tidyverse because of 'select()' conflict
library(tidyverse)	# Data handling
library(patchwork)	# Easy combining of ggplot figures
library(ggtext)	# HTML formatting of axis labels
library(nnet)		# Provides multinom()
library(brms)		# Bayesian models
library(scales)	# Transforms axis scale labels
library(flan)		# Fluctuation test package
library(Cairo)		# Cairo PDF output for Greek/UTF-8 symbols
library(emmeans)
library(afex)
library(car)


## Data Munging ##
######################################################################
## SNPs from Sanger Sequencing
# Reading in, tidying, and assigning factor levels
# First, save the file mutations_cites.csv file from File S1 'mutations' tab.
mutations = read_csv("mutation_sites.csv") %>%
	gather(sample, has.SNP, -c(type:ALT)) %>%
	mutate(has.SNP = as.logical(as.numeric(has.SNP))) %>%
	mutate(plate = substr(sample,1,1)) %>%
	mutate(treatment = recode_factor(plate,
			B = "Glucose 80", C = "Glucose 80",
			A = "Glucose 250", D = "Glucose 250"),
		strain = recode_factor(plate,
			A = "MG1655", C = "MG1655",
			B = "ΔluxS", D = "ΔluxS"),
		strain_treatment = fct_relevel(paste0(strain, ", ", treatment),
			c("MG1655, Glucose 80", "MG1655, Glucose 250",
			"ΔluxS, Glucose 80", "ΔluxS, Glucose 250")),
		mutation = SNP, 
		SNP = gsub("([+-])[CGAT*]*", "indel\\1", SNP),
		SNP = recode_factor(SNP,
			`AT>GC` = "AT>GC", `GC>AT` = "GC>AT",  # Transitions
			`AT>CG` = "AT>CG", `AT>TA` = "AT>TA",  # Transversions
			`GC>CG` = "GC>CG", `GC>TA` = "GC>TA",  # Transversions
			`indel+`="insertion", `indel-`="deletion"), # Indels
		SNP.category = recode_factor(SNP,
			`AT>GC` = "Transitions",   `GC>AT` = "Transitions",
			`AT>CG` = "Transversions", `AT>TA` = "Transversions",
			`GC>CG` = "Transversions", `GC>TA` = "Transversions",
			`insertion` = "Indels",  `deletion` = "Indels")) # Indels


# Retain only rows where sample actually has the SNP after the tidying gather:
SNPs = mutations %>%
	#filter(type == "SNP", has.SNP) %>% # SNVs only
	filter(has.SNP) %>%
	droplevels()
	
######################################################################
## Mutation rate from fluctuation test

# Colony counts on non-selective media (Nt)
Nt_raw = read_csv("File S3 - Nt counts.csv")

# Here comes a bodge... why can't gather() do two columns at once??
Nt_density = Nt_raw %>%
	select(strain_treatment, plate_ID, starts_with("density")) %>%
	gather(key = "key", value = "density_ml", -strain_treatment, -plate_ID) %>%
	mutate(rep = gsub("[A-Za-z_]","", key)) %>%
	select(-key)
	
Nt_Nt = Nt_raw %>%
	select(strain_treatment, plate_ID, starts_with("Nt")) %>%
	gather(key = "key", value = "Nt", -strain_treatment, -plate_ID) %>%
	mutate(rep = gsub("[A-Za-z_]","", key)) %>%
	select(-key)
	
Nt_data = full_join(Nt_density, Nt_Nt)

rm(Nt_density, Nt_Nt, Nt_raw)
# End bodge
	
Nt_summary = Nt_data %>%
	group_by(strain_treatment) %>%
	summarise(mfn = mean(Nt), cvfn = sd(Nt)/mean(Nt),
			density.mean = mean(density_ml),
			density.se = sd(density_ml)/sqrt(length(density_ml)))

# Mutant counts
# NB: wells with too many resistant mutants to count replaced by max(count) + 1

# Estimate the mutation rate, its sd (for plotting) and 95% CI (for ratio testing)
mcdata = read_csv("File S2 - Mutant counts.csv") %>%
	gather(key = strain_treatment, value = mc) %>%
	group_by(strain_treatment) %>%
	nest() %>%
	ungroup() %>%
	left_join(., Nt_summary) %>%
		mutate(mu = pmap(.l = list(data, mfn, cvfn),
			function(a, b, c) mutestim(mc = a$mc, mfn = b, cvfn = c))) %>%
		mutate(mu.test = pmap(.l = list(data, mfn, cvfn),
			function(a, b, c) flan.test(mc = a$mc, mfn = b, cvfn = c))) %>%
		mutate(mutprob.l95 = map_dbl(mu.test, ~.$conf.int[[1,1]]),
			mutprob.u95 = map_dbl(mu.test, ~.$conf.int[[2,1]])) %>%
	separate(strain_treatment,
		into = c("strain", "treatment"), remove = F, sep = ", ") %>%
	mutate(strain_treatment = fct_relevel(strain_treatment,
			c("MG1655, Glucose 80",
			"MG1655, Glucose 250",
			"ΔluxS, Glucose 80",
			"ΔluxS, Glucose 250")),
		treatment = fct_relevel(treatment,
			c("Glucose 80", "Glucose 250")),
		strain = fct_relevel(strain, c("MG1655","ΔluxS"))) %>%
	unnest_wider(mu)

# Use the estimates to calculate a point estimate for the ratio, according to Eqn 1-3 of the supplementary info

muratios = mcdata %>%
	select(strain, treatment, mutprob, sd.mutprob) %>%
	pivot_wider(names_from = treatment, values_from = c(mutprob, sd.mutprob)) %>%
	mutate(ratio.mean = `mutprob_Glucose 80`/`mutprob_Glucose 250`) %>%
	mutate(ratio.sigma2 = 
		(1/`mutprob_Glucose 250`^2)*
		(`sd.mutprob_Glucose 80`^2+
			(`mutprob_Glucose 80`^2/`mutprob_Glucose 250`^2)*
			`sd.mutprob_Glucose 250`^2)) %>%
	select(strain, starts_with("ratio"))

urhatMG = muratios$ratio.mean[muratios$strain == "MG1655"]
urhatLuxS = muratios$ratio.mean[muratios$strain == "ΔluxS"]
sigma2hatMG = muratios$ratio.sigma2[muratios$strain == "MG1655"]
sigma2hatLuxS = muratios$ratio.sigma2[muratios$strain == "ΔluxS"]

	
ratio.zstat = (urhatMG - urhatLuxS)/(sqrt(sigma2hatMG + sigma2hatLuxS))
ratio.pvalue = 1-pnorm(ratio.zstat) # very small


## Models ##
######################################################################
# Model M1: Categorical model treating each SNP as separate \

m1_1 = multinom(SNP ~ strain*treatment, data = SNPs)
m1_2 = multinom(SNP ~ strain+treatment, data = SNPs)
m1_3 = multinom(SNP ~ treatment, data = SNPs)
m1_4 = multinom(SNP ~ strain, data = SNPs)
m1_5 = multinom(SNP ~ 1, data = SNPs)

modelM1a.t = anova(m1_1, m1_2, m1_3, m1_5)
modelM1b.t = anova(m1_1, m1_2, m1_4, m1_5)
modelM1.AIC = AIC(m1_1, m1_2, m1_3, m1_4, m1_5)

set_sum_contrasts() # use sum coding, necessary to make type III LR tests valid
Anova(m1_1,type="II")


# Model M2: Logistic model on each SNP separately
SNPs.indel = SNPs %>%
	mutate(type = type!="SNP")

mINDEL = list(mINDEL0 = glm(type ~ treatment*strain, data = SNPs.indel,
		 family = binomial(link = "logit")),
	mINDEL1 = glm(type ~ treatment+strain, data = SNPs.indel,
		 family = binomial(link = "logit")),
	mINDEL2 = glm(type ~ strain, data = SNPs.indel,
		 family = binomial(link = "logit")),
	mINDEL3 = glm(type ~ treatment, data = SNPs.indel,
		 family = binomial(link = "logit")),
	mINDEL4 = glm(type ~ 1, data = SNPs.indel,
		 family = binomial(link = "logit"))
)


SNPs.bino = SNPs %>%
	dplyr::select(sample, strain, treatment, SNP, has.SNP) %>%
	pivot_wider(names_from = SNP, values_from = has.SNP, values_fill = FALSE)

	
#GC>AT nothing significant
SNPs.bino = SNPs %>%
	mutate(SNP = recode_factor(SNP, `GC>AT`="GC>AT", .default = "other"))

mGCAT = list(mGCAT0 = glm(SNP ~ treatment*strain, data = SNPs.bino,
		 family = binomial(link = "logit")),
	mGCAT1 = glm(SNP ~ treatment+strain, data = SNPs.bino,
		 family = binomial(link = "logit")),
	mGCAT2 = glm(SNP ~ strain, data = SNPs.bino,
		 family = binomial(link = "logit")),
	mGCAT3 = glm(SNP ~ treatment, data = SNPs.bino,
		 family = binomial(link = "logit")),
	mGCAT4 = glm(SNP ~ 1, data = SNPs.bino,
		 family = binomial(link = "logit"))
)

#AT>GC significant effect of treatment, not strain
SNPs.bino = SNPs %>%
	mutate(SNP = recode_factor(SNP, `AT>GC`="AT>GC", .default = "other"))
mATGC = list(mATGC0 = glm(SNP ~ treatment*strain, data = SNPs.bino,
		 family = binomial(link = "logit")),
	mATGC1 = glm(SNP ~ treatment+strain, data = SNPs.bino,
		 family = binomial(link = "logit")),
	mATGC2 = glm(SNP ~ strain, data = SNPs.bino,
		 family = binomial(link = "logit")),
	mATGC3 = glm(SNP ~ treatment, data = SNPs.bino,
		 family = binomial(link = "logit")),
	mATGC4 = glm(SNP ~ 1, data = SNPs.bino,
		 family = binomial(link = "logit"))
)

#ATCG
SNPs.bino = SNPs %>%
	mutate(SNP = recode_factor(SNP, `AT>CG`="AT>CG", .default = "other"))
mATCG = list(mATCG0 = glm(SNP ~ treatment*strain, data = SNPs.bino,
		 family = binomial(link = "logit")),
	mATCG1 = glm(SNP ~ treatment+strain, data = SNPs.bino,
		 family = binomial(link = "logit")),
	mATCG2 = glm(SNP ~ strain, data = SNPs.bino,
		 family = binomial(link = "logit")),
	mATCG3 = glm(SNP ~ treatment, data = SNPs.bino,
		 family = binomial(link = "logit")),
	mATCG4 = glm(SNP ~ 1, data = SNPs.bino,
		 family = binomial(link = "logit"))
)

# GCTA
SNPs.bino = SNPs %>%
	mutate(SNP = recode_factor(SNP, `GC>TA`="GC>TA", .default = "other"))
mGCTA = list(mGCTA0 = glm(SNP ~ treatment*strain, data = SNPs.bino,
		 family = binomial(link = "logit")),
	mGCTA1 = glm(SNP ~ treatment+strain, data = SNPs.bino,
		 family = binomial(link = "logit")),
	mGCTA2 = glm(SNP ~ strain, data = SNPs.bino,
		 family = binomial(link = "logit")),
	mGCTA3 = glm(SNP ~ treatment, data = SNPs.bino,
		 family = binomial(link = "logit")),
	mGCTA4 = glm(SNP ~ 1, data = SNPs.bino,
		 family = binomial(link = "logit"))
)

#mGCCG
SNPs.bino = SNPs %>%
	mutate(SNP = recode_factor(SNP, `GC>CG`="GC>CG", .default = "other"))
mGCCG = list(mGCCG0 = glm(SNP ~ treatment*strain, data = SNPs.bino,
		 family = binomial(link = "logit")),
	mGCCG1 = glm(SNP ~ treatment+strain, data = SNPs.bino,
		 family = binomial(link = "logit")),
	mGCCG2 = glm(SNP ~ strain, data = SNPs.bino,
		 family = binomial(link = "logit")),
	mGCCG3 = glm(SNP ~ treatment, data = SNPs.bino,
		 family = binomial(link = "logit")),
	mGCCG4 = glm(SNP ~ 1, data = SNPs.bino,
		 family = binomial(link = "logit"))
)

#ATTA
SNPs.bino = SNPs %>%
	mutate(SNP = recode_factor(SNP, `AT>TA`="AT>TA", .default = "other"))
mATTA = list(mATTA0 = glm(SNP ~ treatment*strain, data = SNPs.bino,
		 family = binomial(link = "logit")),
	mATTA1 = glm(SNP ~ treatment+strain, data = SNPs.bino,
		 family = binomial(link = "logit")),
	mATTA2 = glm(SNP ~ strain, data = SNPs.bino,
		 family = binomial(link = "logit")),
	mATTA3 = glm(SNP ~ treatment, data = SNPs.bino,
		 family = binomial(link = "logit")),
	mATTA4 = glm(SNP ~ 1, data = SNPs.bino,
		 family = binomial(link = "logit"))
)

# Type II ANOVA
## Models with significant terms:
mATGC[[1]] %>% Anova(type=2)
mGCTA[[1]] %>% Anova(type=2)

## Models with non-significant terms
mGCAT[[1]] %>% Anova(type=2)
mATCG[[1]] %>% Anova(type=2)
mATTA[[1]] %>% Anova(type=2)
mGCCG[[1]] %>% Anova(type=2)

## Plots ##
######################################################################

# Plotting colours
cols = c("#2a859e", "#0a4454", "#d15938", "#87280e")
cols2 = c("#2a859e", "#0a4454", "#d15938", "#87280e", "#87460d", "#d17f38")
cols.s = c("#bc481a","#f27b24", "#143d3c", "#0c6860", "#00a395", "#35d6c8")
cols.t = c("#e55e5e", "#bf0000", "#5b0000", "#44d0ff", "#007ca5", "#0a2b35")

######################################################################
# Figure 1a: Mutation rates
theme_set(theme_bw())
theme_update(text = element_text(size=12),
	panel.grid.major = element_blank(),
	panel.grid.minor = element_blank(),
	strip.background = element_blank())

mu.p = mcdata %>% ggplot(aes(x = density.mean/10^8,
			y = mutprob*10^8, colour = strain_treatment)) +
	geom_point(size = 3) +
	geom_errorbar(aes(#x = treatment, y = mutprob*10^8,
		ymin = mutprob*10^8 - sd.mutprob*10^8,
		ymax = mutprob*10^8 + sd.mutprob*10^8),
		width = 0) +
	geom_errorbarh(aes(
		xmin = density.mean/10^8 - density.se/10^8,
		xmax = density.mean/10^8 + density.se/10^8,
		height = 0)) +
	scale_x_log10(breaks = c(0.5,1,2), limits = c(0.5,4)) +
	scale_y_log10(breaks = seq(from = 0.75, to = 2, by = 0.25)) +
	scale_colour_manual(name = "Strain, treatment",
		values = cols.t[c(1,2,4,5)]) +
	labs(x = "Density (×10<sup>8</sup> cells)",
		y ="Mutation rate<br>(×10<sup>-8</sup> per generation)") +
	theme(axis.title.x = element_markdown(), axis.title.y = element_markdown())
		
#cairo_pdf("figure1a.pdf", width=4, height=2)
#	mu.p
#dev.off()

# Figure 1b: Spectrum
# Pyramid plot - strain
# Reflect the frequencies around the x axis to compare between strains/treatments
# Frequencies also adjusted by proportion of GC sites that are known rifR sites
theme_set(theme_bw())
theme_update(text = element_text(size=12),
	panel.grid.major = element_blank(),
	panel.grid.minor = element_blank(),
	strip.background = element_blank())

pyramid_data_strain = SNPs %>%
	group_by(strain, treatment, strain_treatment, SNP, SNP.category) %>%
	summarise(count = n()) %>%
	ungroup() %>%
	group_by(strain, treatment, strain_treatment) %>%
	mutate(freq = count / sum(count)) %>%
	mutate(freq = ifelse(treatment=="Glucose 250", freq*-1, freq)) %>%
	mutate(freq.adj = ifelse(grepl("GC>", SNP)&treatment!="MA",
		(freq/0.605)/2, (freq/(1-0.605))/2)) %>%
	ungroup()

pyramid_plot_strain = pyramid_data_strain %>%
	ggplot(aes(x=SNP, y = freq, fill = strain_treatment), width = 0.4, colour = "black") +
	geom_bar(data = subset(pyramid_data_strain, strain=="MG1655" ), stat = "identity", position="dodge") +
	geom_bar(data = subset(pyramid_data_strain, strain!="MG1655"), stat = "identity", position="dodge") +
	scale_y_continuous(breaks = c(-0.6, -0.4, -0.2, 0, .2, .4,.6),
		labels = c(0.6,0.4,0.2,0,0.2,0.4,0.6), limits =c (-0.5,0.5)) +
		facet_grid(strain~SNP.category, scales = "free", space = "free_x") +
	scale_fill_manual(values =  cols.t[c(2,1,5,4)], name = "Treatment",  guide = 
		guide_legend(reverse = F)) +
	geom_hline(yintercept = 0, size = 0.25) +
	labs(x = "SNP", y = "Frequency") +
	theme(axis.text.x = element_text(angle = 45, hjust = 1))
pyramid_plot_strain

#ggsave("figure1b.pdf", pyramid_plot_strain, width = 6, height = 4, device=cairo_pdf)

## Table S1 - Generate LaTeX code ##
######################################################################
mutations %>%
	filter(has.SNP) %>%
	select(strain_treatment, plate, sample, sitebp, 
		type, mutation, SNP, REF, ALT, aachange) %>%
	arrange(sitebp) %>%
	group_by(strain_treatment, sitebp, SNP, mutation, aachange) %>%
	summarise(count = n()) %>%
	pivot_wider(names_from = strain_treatment, values_from = count, values_fill=0) %>%
	arrange(sitebp) %>%
	mutate(mutation = gsub("\\*","", mutation)) %>%
	mutate(length = nchar(mutation)-1) %>%
	mutate(sitebp = ifelse(SNP%in%c("insertion","deletion"),
		paste(sitebp, sitebp+length-1, sep="-"), paste(sitebp))) %>%
	ungroup() %>%
	select(-c(SNP, length)) %>%
	xtable::xtable(digits = 0) %>%
	print(include.rownames = F)
```
